# Single neurons in the human substantia nigra encode social learning signals

**DOI:** 10.64898/2026.09.18.752742

**Authors:** Arianna N. Davis, Ofer Perl, Brian H. Kopell, Matthew Heflin, Qi Xiu Fu, Salman E. Qasim, Zarghona Imtiaz, Ayaka Kato, Soojung Na, Navid Mikail, Helen S. Mayberg, P. Read Montague, Ignacio Saez, Xiaosi Gu

## Abstract

Humans are remarkable at adapting to changing norms, yet we know little about how the human brain encodes these social signals at the single neuron level. Here, we recorded the activity of single neurons in the substantia nigra and globus pallidus in neurosurgical participants as they played an iterative ultimatum game versus human and computer avatars. Using computational modeling of behavior, we found that putative dopaminergic units in the substantia nigra tracked the valence of norm prediction errors, with greater encoding during gameplay with human versus computer avatars. Analogous analyses in the globus pallidus, a nucleus adjacent to the substantia nigra, found no significant effect. Our results identify a novel role for the substantia nigra in encoding social learning signals.

**AUTHOR SUMMARY:** An important skill within social functioning is the ability to learn and adapt to changing social norms, or rules of behavior among social groups. While computational modeling of behavior has advanced our understanding of how humans learn social norms, little is known about what neural mechanisms support this process. Here, we leverage computational modeling of behavior in conjunction with human intracranial recording to investigate how the human brain encodes norm learning signals at the single neuron level. We recorded the activity of single neurons in the human substantia nigra pars reticulata (SNr) and globus pallidus internus (GPi) in neurosurgical participants as they played a social exchange game with human and computer avatars. We estimated trial-by-trial norm prediction errors (nPEs), or mentally computed errors in social prediction, and revealed that putative dopaminergic SNr neurons encoded the valence of nPEs, with greater encoding observed during gameplay with human versus computer avatar partners. This finding was not observed in the GPi. Together, our results expand upon existing knowledge about the SNr by identifying a novel role for SNr neurons in social norm learning.

## INTRODUCTION

Humans are inherently social creatures.^1–8^ One key inquiry into the social nature of human behavior has centered around our perception of and response to social norms defined as culturally accepted rules or patterns of behavior among social groups.^9–12^ Previous work has conceptualized norms as relatively stable social perceptions. More recently, however, computational modeling of behavior has started to elucidate the dynamic and flexible nature in the human perception of norms by showing that humans can adjust to changing social environments based on norm prediction errors (nPEs), defined as the difference between a social signal or offering and one’s internal expectation.^13–16^ Neuroimaging and lesion evidence has shown a role of anterior insula and ventral striatum in tracking nPEs. ^13,16^ Yet, due to certain limitations of these methods (e.g. low spatiotemporal resolution), how deep nuclei in the mesolimbic system, such as the substantia nigra (SN), may be involved in norm adaptation remains elusive.

Much of what we know about SN function has been established by animal work. For example, it has been well documented that dopaminergic (DA) projections from the SN play a key role in reward learning.^17–21^ While direct study of the human SN and broader DA system is more challenging with standard non-invasive techniques such as BOLD fMRI, recent studies have used invasive recording methods (i.e. single-unit recording) to characterize a role for the SN in reinforcement learning.^22,23^ Note, the SN is comprised of the SN pars compacta (SNc), which contains primarily DA neurons with fewer GABAergic neurons, and the SN pars reticulata (SNr), which contains primarily GABAergic neurons with comparatively fewer DA neurons.^24–26^ Given that DA and GABAergic neuronal populations are not perfectly separated according to anatomical landmarks in the SN,^27^ human single-unit studies often isolate putative DA neuron activity according to characteristic properties of DA waveforms recorded by microelectrodes without specifying recording sites as SNr or SNc.^22,27^ In doing so, such work has demonstrated that putative DA neurons in the human SN (possibly SNr and/or SNc) exhibit increased firing rates after unexpected financial gains than unexpected losses.^22^ Conversely, phasic microstimulation of these cells inhibits learning.^23^ These findings are consistent with literature implicating the SN in reward learning^28^ and DA units in reward prediction error encoding.^29–31^

In addition to this central role in reward learning, DA activity has also been shown to be important for social behavior in human^32–34^ and rodent^35,36^ studies. Human pharmacological studies have demonstrated that DA activity modulates altruism^33^ and egalitarian behavior.^34^ Rodent studies have demonstrated that DA activity encodes information about unfamiliar conspecifics, akin to a social prediction error.^35^ Conversely, DA dysfunction causes behavioral inflexibility, increased reactivity, and aggression.^36^ One recent study used fast-scan cyclic voltammetry in humans as they played an ultimatum game to directly examine the role of DA in social behavior, specifically in the SNr.^32^ This study revealed that extracellular dopamine concentrations in the SNr track differences between previous and current offers, with higher overall dopamine concentrations observed when subjects play with human as opposed to computer avatars.^32^ However, the role of individual SN neurons in human social behavior remains unknown. More generally, it would be valuable to formally relate the body of work demonstrating a role for the DA system in reward learning and the work demonstrating its role in social behavior at the single neuron level.

Here, we leverage computational modeling of behavior and human single-unit recordings to determine the computational role of SNr neurons in social norm behavior. Participants undergoing deep brain stimulation (DBS) device implantation surgery for Parkinson’s disease (PD) underwent single-unit recording from the SNr or globus pallidus internus (GPi) while playing an iterated ultimatum game (UG), a behavioral paradigm that has been previously used to probe response to fairness^37,38^ as well as norm adaptation.^14^ Overall, participants chose to accept or reject variable offers from human or computer partners. As previously described, participants often rejected disadvantageous offers but were able to adapt to counterpart offers (i.e. social norm) over time. Computational modeling of behavior revealed that participants expected higher offers when playing with human versus computer partners, suggesting fairness expectations depended on social partner identity. Importantly, the valence of nPEs, which signal the direction of deviations between expected and received offers, was encoded in the activity of putative DA SNr neurons, with greater encoding during gameplay with human versus computer avatars. An analogous control experiment in the GPi found no significant effect of nPE valence on firing rates. These results demonstrate that norm learning signals drive SNr activity during social interactions and provide a neurocomputational characterization of this activity at the single neuron level.

## RESULTS

### Experimental framework

We collected 27 single-unit recording sessions from the SNr in 19 participants with PD (16 men, 3 women, aged 65 ± 1.93 years (mean ± SEM), **S1 Table**) who underwent DBS implantation surgery in the subthalamic nucleus (STN) (**Figure 1A**). During single-unit recordings, participants played an iterated version of UG in which offers dynamically changed over time. On each trial, participants were presented with a proposed split of $20 by a fictional “human” or “computer” partner, which they could accept or reject (**Figure 1B**). If the participant accepted the offer, each party received the proposed split. If the participant rejected the offer, neither party received a reward. Offers were pseudorandomized and normally distributed, ranging from $1 to $9 with a mean offer size of $5. This game also included self-reported mood ratings after a random 33% of trials in each block. Before behavioral analyses, sessions with flat behavior were excluded in order to protect data quality from inattentiveness, drowsiness, or poor comprehension. Flat sessions were defined as those in which less than 2 offers were accepted or rejected in the entirety of a human or computer block, and those in which a logistic fit of choices versus offer size had a non-positive slope. In the SNr cohort, 3 human avatar blocks and 3 computer avatar sessions were excluded. For further information, see **Materials and Methods.**

**Figure 1.**
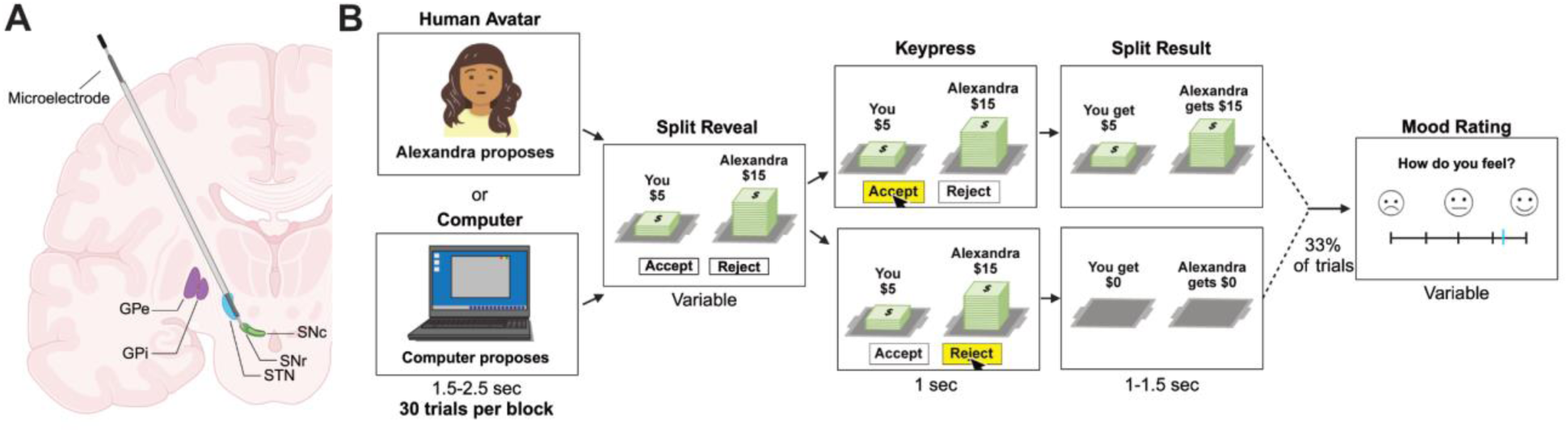
Experimental framework. **(A)** Intraoperative targeting of the microelectrode in the substantia nigra pars reticulata (SNr). Microelectrodes were positioned ventral to the clinical target, the subthalamic nucleus (STN), for placement in the recording target, the SNr. The position of the microelectrode and relative anatomical locations of the brain regions along its trajectory were identified in real time by surgical mapping, visualization of neuronal spikes, and audition of neural firing with an Alpha Omega electrophysiological recording system, such that each region could be identified by its unique signature of neural activity. **(B)** Ultimatum game. Participants played one of two versions of the ultimatum game. In one version, human avatar partners repeatedly present participants with proposed splits of $20, which the participants accept or reject. If the participant accepts, each party receives the proposed split. If the participant rejects, each party receives nothing. Each offer is presented by a new partner. All offers are less than $9. A second version of the ultimatum game includes not only a human block but also a block of trials with “computer” partners, as well as a mood rating after a random 33% of choice trials. Brain slice and computer icon created in BioRender. Neal, A. (2027) https://BioRender.com/bpwujj1

### Ultimatum game behavior

Previous work has found that healthy humans are more likely to reject unfair offers,^14,37,38^ especially from human avatar partners.^37^ Therefore, we first sought to examine how our participants’ choices differed as a function of the fairness of the offers and the decision context (human vs. computer condition) by running a mixed-effects model, whereby binary choices (reject = 0, accept = 1) were predicted by offer size (i.e. $1, 2, etc.), condition (human = 1, computer = 2), and the interaction between these terms, with random effects for participant and for session within participant. Only the impact of offer size significantly impacted choices (offer, \*\*\**p* = 8.2e-31, **β** ± SE = -0.85 ± 0.071; OR = 0.43, 95% CI [0.37, 0.49]; condition, *p* = 0.15, **β** ± SE = -0.93 ± 0.65, OR = 0.40, 95% CI [0.11, 1.41]; offer x condition, *p* = 0.21, **β** ± SE = 0.14 ± 0.12, OR = 1.15, 95% CI [0.92, 1.45]), indicating that participants’ rejection rates decreased as a function of increasing offer size across conditions (**Figure 2A**). A mixed-effects model predicting overall rejection rate by condition with the same random effects demonstrated that overall rejection rate difference between conditions was not significant (condition, *p* = 0.46, **β** ± SEM = -0.14 ± 0.18; OR = 0.87, 95% CI [0.61, 1.25]) (**Figure 2B**). Analogous mixed-effects models demonstrated that overall reaction times (condition, *p* = 0.29, **β** ± SEM = 0.43 ± 0.41; OR = 1.53, 95% CI [0.69, 3.41]; **S1A Figure**) and mood ratings (condition, *p* = 0.68, **β** ± SEM = 1.00 ± 2.41; OR = 2.72, 95% CI [0.023, 320.83]; **S1B Figure**) did not differ significantly between conditions. However, a mixed-effects model predicting z-scored mood rating by the immediately preceding choice, condition, and the interaction between these terms with participant and session random effects demonstrated that z-scored mood ratings were higher after accepted trials than rejected trials in both the human and computer conditions (choice, \**p* = 0.024, **β** ± SEM = 0.50 ± 0.22; OR = 1.64, 95% CI [1.07, 2.53]; condition, *p* = 0.81, **β** ± SEM = -0.049 ± 0.20; OR = 0.95, 95% CI [0.64, 1.41]; condition x choice, *p* = 0.63, **β** ± SEM = 0.15 ± 0.31; OR = 1.16, 95% CI [0.62, 2.17]) **(Figure 2C**).

**Figure 2.**
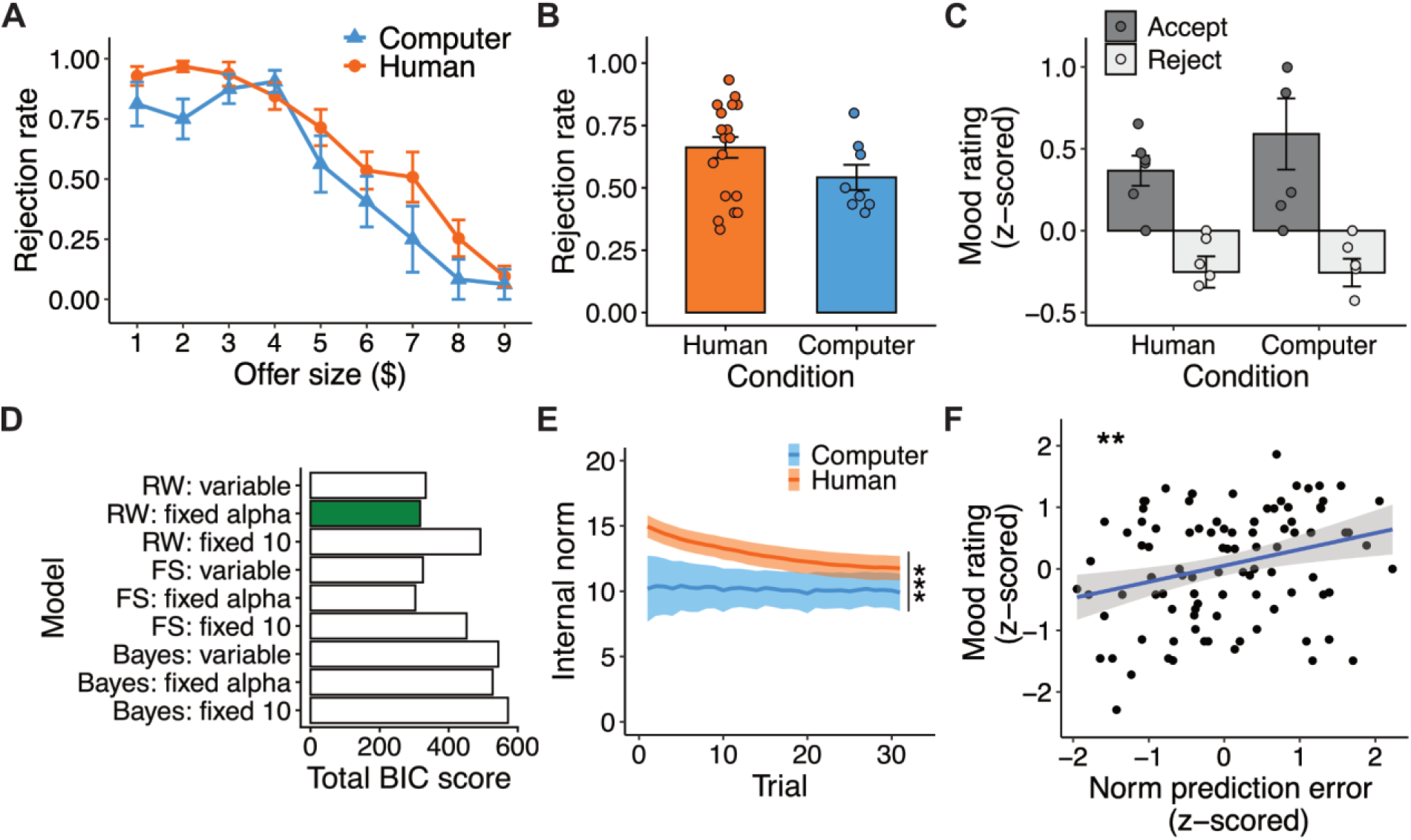
Ultimatum game behavior. **(A)** Rejection rate by offer size and **(B)** overall rejection rate in the computer (blue) and human (orange) conditions. **(C)** Mood rating (z-scored) versus choice (accept/reject) in the human and computer conditions. **(D)** Comparison of norm adaptation models by total Bayesian information criterion (BIC) score for all sessions and participants. The model with the lowest total BIC score has the best fit (green). **(E)** Internal norm by trial in the computer and human conditions. Internal norms were calculated using the best performing norm adaptation model. **(F)** Norm prediction error (z-scored) versus mood rating (z-scored) for all sessions and participants. Bars, points, and lines represent the mean. Error bars represent standard error of the mean. \*\**p* < 0.01; \*\*\**p* < 0.001 by two-tailed Wilcoxon rank sum tests **(C)**, general linear regression **(E)**, and Pearson’s product-moment correlation test **(F)**. *n* = 21 blocks of gameplay with human avatars and 8 blocks of gameplay with the computer avatar for (A), (B), (D), and (E). *n* = 6 blocks of gameplay with human avatars and 5 blocks of gameplay with the computer avatar for (C) and (F).

Together, these results indicate that participants were averse to low offers and experienced negative momentary mood states after rejecting offers. These findings suggest that despite having Parkinson’s, and even without monetary incentive, our participants behaved in a way that was largely similar to previous findings regarding the ultimatum game in healthy, younger adults.^14,16,37,38^

### Modeling norm adaptation

Next, we carried out computational modeling of behavior to characterize rejection behavior and the computational dynamics underlying subjective norm updating. All modeling details are listed in **Materials and Methods**. Briefly, similar to our previous work, we constructed the subjective utility U(t) at each trial as the difference between the offer received and the individual’s sensitivity to norm violation.^14,15^ In the model, *f* represents the individual’s internal norm, and alpha (α ∈ [0,1]) represents the participant’s sensitivity to norm violation (i.e. perceived differences between the internal norm and the received offer), with higher α values representing greater aversion to unfairness. Subjectivity utility U(t) is related to choice probability through a softmax function.^14^

We compared three classes of models that varied in their assumptions about how internal norms are updated: Rescorla-Wagner (RW) models^14,39,40^ in which norms are updated using temporal difference learning; Bayesian (Bayes) models^13,14^ in which norms are updated following Bayes’ rule; and Fehr-Schmidt (FS) models of inequity aversion^14,15^ in which norms are not updated. Within each class, individual models differed by their assumptions about the initial internal norm (f0), which could be fixed at $10 (fixed f0, corresponding to an equal split) or a variable initial norm expectation that was individually fit to each participant’s behavior (variable f0). In other words, if one player begins the session with a fairness expectation for an initial offer of $9 and another player begins with an expectation for an initial offer of $11, then the variable f0 model should capture each of those initial expectations during model fitting. So, in models with a variable f0, f0 was a free parameter. Additionally, we also considered models with variable f0 in which the sensitivity to unfairness α was fixed to be the group’s average α (fixed α); the group’s average α was obtained by taking the average of estimates for α originally fit as a free parameter. Therefore, we considered a total of 9 models (3 models x 3 initial internal norm/alpha specification). For variable f0 models, f0 and α were both free parameters. For fixed f0 models, α was a free parameter and f0 was fixed. Finally, for fixed α models, f0 was a free parameter and α was fixed.

After fitting models to each participant’s behavioral data, we compared fit quality by examining the resulting minimum Bayesian information criterion (BIC) score at both the individual (**S9 Table2 Dataset**) and group level (**Figure 2D**). The FS: fixed α model produced the best fit, followed by the RW: fixed alpha model (**Figure 2D**).

Additionally, in order to explore whether behavior might be driven by a classic RPE, we compared all 9 models to a Q-learning approach, similar to previous work.^16^ A classic Q-learning algorithm learns an optimal action-value function within an environment without any prior model of the dynamics of that environment. It does so by updating estimates of the value of taking an action in a given state (i.e. Q-value) based on the reward received and the estimated value of the next state. This method adjusts Q-values incrementally as driven by RPEs. RPE-directed updates are made at a learning rate between 0 and 1 after the reward is received and before the next offer is proposed. So, while an nPE describes how much higher or lower an offer is above or below an individual’s model-based norm for fairness, an RPE, on the other hand, represents how much higher or lower an outcome was versus an expectation based on previous events. We compared the fit quality of all 9 models to that of the Q-learning approach by minimum total BIC score. (See **S1 Dataset** and **S2 Table** for all parameter estimates as well as distributions of the envy parameter, respectively, for each participant and each model.) Model comparison revealed that the Q- learning approach performed worse than all 9 models, with the FS: fixed α and RW: fixed α models again producing the best fit (**S2A Figure**; see **S2 Dataset** for individual BIC scores). Furthermore, these results are consistent both across and between human and computer conditions (**S2B-C Figure**). These results suggest that behavior in our task is motivated not by classic Q-learning, but rather by static preferences based on social variables, as in the FS: fixed α model, or by model-driven incorporation of social information into norm expectations over time, as in the RW: fixed α model.

Next, to facilitate better judgment of the strength of evidence for each of the two best-fitting models, we assessed ΔBIC, expected model frequency, and protected exceedance probabilities (S3 Table). These analyses demonstrated that the FS: fixed α model had the lowest BIC with an expected model frequency of 0.435 and a protected exceedance probability of 0.932. The RW: fixed α model had a ΔBIC of 13.707, expected model frequency of 0.174, and protected exceedance probability of 0.043. So, while there is greater evidence in support of the FS: fixed α model overall, there is also evidence of some heterogeneity in best fit between the two candidate models, consistent with the fact that mean BIC scores were comparable at 10.45 and 10.93 for the FS: fixed α and RW: fixed α models, respectively.

We then continued our assessment of these two models by exploring the degree to which each model was identifiable via model recovery (S3 Figure; **S4 Table)**. Model recovery analysis demonstrated a recovery probability of 45.47% for the RW: fixed alpha model and 37.62% for the FS: fixed alpha model, indicating that while these models performed comparably well for best fit, the RW: fixed alpha model demonstrated higher identifiability. Additional recovery at the class-level (FS models, RW models, Bayes models, Q-learning), performed by averaging recovery probabilities across models within each class, further confirmed that the behavioral predictions generated by these computational model classes are distinguishable (S5 Table; S4 Figure). So, out of the two best-fitting models, the RW: fixed α model demonstrated the best identifiability on model recovery.

We next examined whether the RW: fixed α model reproduces the choice patterns observed in our study by conducting both parameter recovery and posterior predictive checks. True (**Table 1**) and simulated parameter values were significantly correlated in this model (**S5 Figure**), indicating good parameter recoverability. Moreover, the RW: fixed *α* model reproduced overall rejection rates in both the human (mean ± SEM: observed: 0.662 ± 0.042; simulated: 0.652 ± 0.045) and computer (observed: 0.542 ± 0.050; simulated: 0.500 ± 0.065) avatar conditions(**S6A Figure).** The RW: fixed *α* model also recapitulated the negative impact of offer size on rejection rate (**S6B Figure)**. Finally, to assess whether the RW: fixed *α* model recapitulates behavior at the individual subject level, individual rejection rates for observed versus simulated sessions were also compared, revealing a strong linear correlation (**S6C Figure**; Pearson’s product-moment correlation, \*\*\**p* = 4.55e-12, r = 0.914, 95% CI 0.823 - 0.959).

**Table 1.**
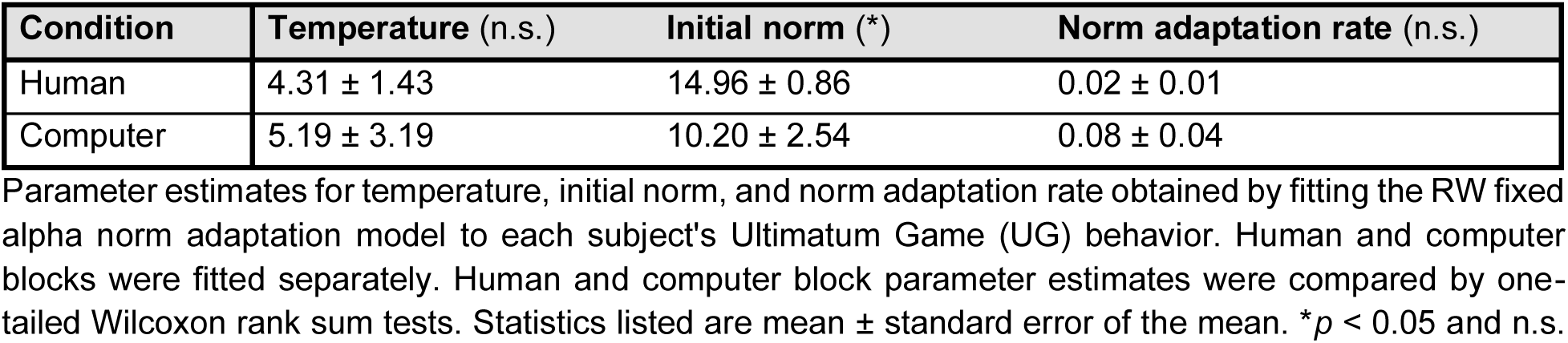
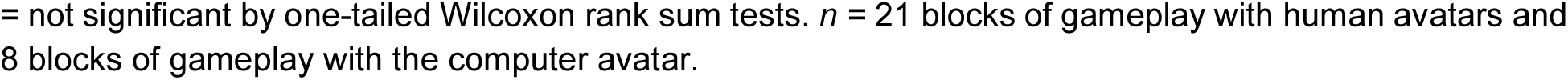
Empirical parameter estimates.

| Condition | Temperature (n.s.) | Initial norm (*) | Norm adaptation rate (n.s.) |
| --- | --- | --- | --- |
| Human | $4.31 \pm 1.43$ | $14.96 \pm 0.86$ | $0.02 \pm 0.01$ |
| Computer | $5.19 \pm 3.19$ | $10.20 \pm 2.54$ | $0.08 \pm 0.04$ |

Parameter estimates for temperature, initial norm, and norm adaptation rate obtained by fitting the RW fixed alpha norm adaptation model to each subject's Ultimatum Game (UG) behavior. Human and computer blocks were fitted separately. Human and computer block parameter estimates were compared by one-tailed Wilcoxon rank sum tests. Statistics listed are mean ± standard error of the mean. $*p < 0.05$ and n.s.
= not significant by one-tailed Wilcoxon rank sum tests. $n = 21$ blocks of gameplay with human avatars and 8 blocks of gameplay with the computer avatar.

Together, the successful model recovery, parameter recovery, and ability of the RW: fixed *α* model to reproduce the behavioral patterns observed in our study at both group and individual levels, providing evidence that this model is suitable for interpreting the mental models driving behavior in our task. Furthermore, these results indicate that temporal-difference learning with fixed unfairness sensitivity best accounts for participants’ incorporation of new information into their norm expectations over time.

Next, we examined the parameter values from the winning RW: fixed α model. We found that both initial and session-wide internal norms (*f*) were higher in the human condition than in the computer condition (**Figure 2E**; **Table 1**). Specifically, a general linear regression predicting internal norm by condition, trial, and the interaction between these terms revealed a significant impact only for condition, whereby internal norms are higher in the human condition (condition, \*\*\**p* = 8.9e-05, **β** ± SEM = 2.5 ± 0.64; trial, *p* = 0.38, **β** ± SEM = -0.027 ± 0.030; trial x condition, *p* = 0.79, **β** ± SEM = 0.0097 ± 0.036). Despite the lack of overall rejection difference between conditions, this model-based analysis suggests that participants had higher expectations for offers from human versus computer avatars.

Because monetary PEs have been shown to impact momentary feelings,^41^ we also examined if this might be the case for nPEs. We calculated nPEs by taking the differences between the current offer size and the preceding internal norm. (For an example of nPE estimation based on choice behavior, see **S7 Figure.)** Positive/negative nPEs indicate offers that are above/below internal norms, respectively. We observed a significant positive correlation between nPEs and the immediately subsequent mood ratings (Pearson’s product-moment correlation, \*\**p* = 0.0012, r = 0.31, 95% CI 0.13 - 0.47) (**Figure 2F**). A mixed-effects model predicting mood ratings by nPEs and condition with a random effect for session resulted in a significant fixed effect only for nPE (nPE, \*\**p* = 0.0013, **β** ± SEM = 0.31 ± 0.09; condition, *p* = 0.96, **β** ± SEM = -0.0089 ± 0.18; OR = 1.36, 95% CI [1.13, 1.63]). Additionally, to determine whether the nPE-mood relationship persists after accounting for choice, we constructed a mixed effects model predicting z-scored mood by z-scored nPE and choice with a random effect for session. The relationship between nPE and mood remained significant after including choice as a fixed effect (nPE, \**p* = 0.035, **β** ± SEM = 0.24 ± 0.11; OR = 1.27, 95% CI [1.02, 1.59]), while choice did not exhibit a significant effect on mood (choice, *p* = 0.32, **β** ± SEM = 0.23 ± 0.22; OR = 1.25, 95% CI [0.81, 1.95]). So, while condition did not impact mood ratings, nPE influenced mood such that negative nPEs decreased mood ratings, consistent with previous work.^41^

### Putative DA neurons in the SNr encode social prediction error valence

Our main goal was to examine the computational role of SNr neurons in norm adaptation. Therefore, we next derived and analyzed single-unit spiking activity from SNr microelectrode recordings. We identified 69 putative neuronal clusters from 27 SNr microelectrode recordings (mean ± SEM clusters per recording: 2.6 ± 0.11) using the OSORT offline sorting algorithm.^42^ Clusters refer to groups of spikes determined by the offline sorting algorithm to be originating from a single source (i.e. putative neuron). Of those 69 clusters, we classified 40 units as putative DA neurons based on mean baseline firing rate (≤15 Hz) and waveform morphology (duration >0.8 ms, biphasic) criteria established by previous work^27^ in human single-unit recording from the SN (mean ± SEM putative DA units per recording: 1.5 ± 0.18) (**Figure 3A)**. Patient-specific stereotactic coordinates for each recorded putative DA neuron are listed in **S6 Table**. Waveform widths and average firing rates for all putative DA neurons are listed in **S7 Table**. Number of putative DA neurons recorded per participant are listed in **S8 Table**. The waveforms for all 40 putative DA units and the mean waveform for these units are depicted in **Figure 3B**. For each unit, we smoothed spike trains into continuous firing rates and z-scored those firing rates to account for variability in firing rates across units in population analyses (**S8 Figure**). We then examined z-scored firing rates of putative DA units at a behaviorally relevant time point in the task, namely around split reveal (the point at which the offers are presented).

**Figure 3.**
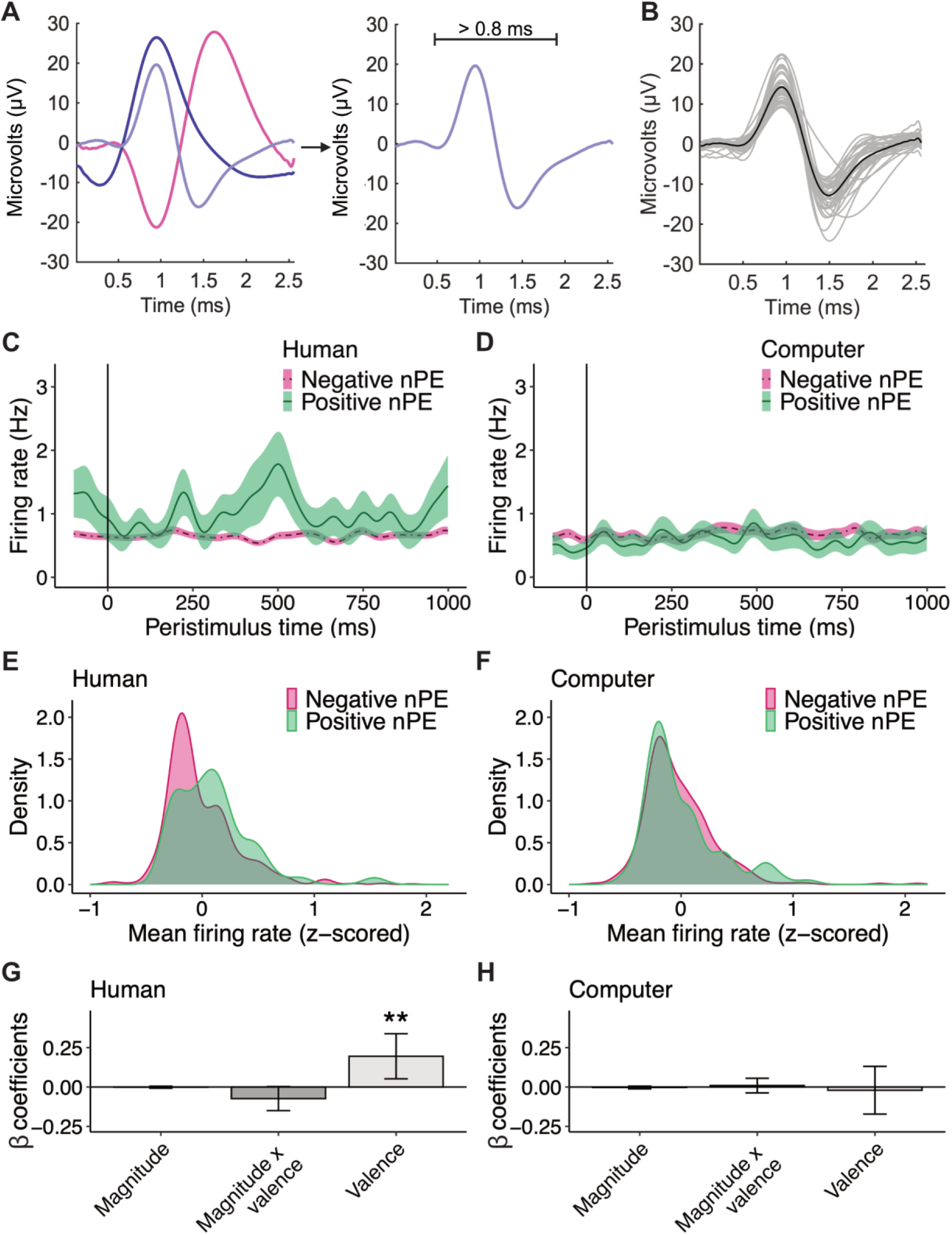
Putative DA unit activity encodes social prediction error valence. **(A)** Three example putative units identified using an offline sorting algorithm and one putative DA unit (light purple) phenotyped by average waveform width >0.8 ms, biphasic morphology, and firing rate ≤15 Hz. **(B)** Average waveform (black) for SNr putative DA units (grey). Mean raw firing rates of SNr putative DA units during the 1000 ms after split reveal for trials producing positive (green, solid) versus negative (magenta, dashed) nPEs in the **(C)** human and **(D)** computer conditions. Split reveal occurs at time 0 ms. Probability density functions of mean firing rates of SNr putative DA units during the 1000 ms after split reveal for trials producing positive (green) versus negative (magenta) norm prediction errors (nPEs) in the **(E)** human and **(F)** computer conditions. β coefficients for the mixed-effects models predicting mean firing rate of SNr putative DA units during the 1000 ms after split reveal in the **(G)** human and **(H)** computer conditions by nPE valence, nPE magnitude, and the interaction terms for these regressors with a random effect for unit. Error bars represent standard error of the mean. \*\**p* < 0.01 by mixed-effects model. *n* = 40 putative DA neurons total for (B). *n* = 31 putative DA neurons in the human condition for (C), (E), and (G). *n* = 16 putative DA neurons in the computer condition for (D), (F), and (H).

We hypothesized that the activity of putative DA neurons may encode certain attributes of nPEs, in a similar way to their known role in representing non-social learning signals.^29–31^ To test this, we first compared mean firing rates of putative DA units during the 1000 ms after split reveal for trials resulting in positive versus negative nPEs (above and below the internal norm, respectively) in the human and computer conditions. See absolute raw continuous firing rates (before z-scoring) in **Figures 3C-D**. See probability density plots of the mean z-scored firing rates during the 1000 ms after split reveal in **Figures 3E-F**. We observed that, in the human condition, putative DA units transiently increased their spiking activity in positive nPE trials higher than in negative nPE trials (**Figure 3C**; **Figure 3E**). This increase was attenuated in the computer condition (**Figure 3D**; **Figure 3F**). To test the significance of the observed impact of nPE valence on firing rate, and also to examine whether these units might also encode nPE magnitude, we constructed a mixed-effects model predicting DA unit activity during the 1000 ms after split reveal by nPE valence (1 = negative, 2 = positive), nPE magnitude, and the interactions between these terms with a random effect for unit in each condition (**Figure 3G-H**; **Table 2**). The fixed effect for nPE valence in the human condition was the only significant predictor of firing rate (nPE valence, \*\**p* = 0.008, **β** ± SEM = 0.19 ± 7.3e-02, 95% CI [0.05, 0.34]; **Figure 3G**; see **Table 2** for other estimates).

**Table 2.** Summary of mixed-effects models predicting substantia nigra pars reticulata (SNr) putative dopaminergic (DA) unit activity at split reveal.

| Fixed effect | Estimate | Std. error | t-value | p-value |
| --- | --- | --- | --- | --- |
| **Human** |  |  |  |  |
| Intercept | -0.004703 | 0.027146 | -0.173 | 0.86280 |
| nPE magnitude | -0.001546 | 0.003287 | -0.470 | 0.63876 |
| **nPE valence \*\*** | 0.194946 | 0.073006 | 2.670 | **0.00771** |
| nPE magnitude x nPE valence | -0.073754 | 0.038881 | -1.897 | 0.05815 |
| **Computer** |  |  |  |  |
| Intercept | -0.003115 | 0.031721 | -0.098 | 0.922 |
| nPE magnitude | -0.002899 | 0.004358 | -0.665 | 0.508 |
| nPE valence | -0.020615 | 0.077226 | -0.267 | 0.790 |
| nPE magnitude x nPE valence | 0.008925 | 0.023744 | 0.376 | 0.707 |

Summary of mixed-effects models in which fixed effects norm prediction error (nPE) magnitude, nPE valence, and their respective interaction terms predicted group activity of SNr putative DA units during the 1000 ms after split reveal in the human or computer condition. $**p < 0.01$ by mixed-effects model. Std. Error = standard error. $n = 31$ putative DA neurons in the human condition and 16 putative DA neurons in the computer condition.

To formally test how the nPE valence effect differs across conditions, we constructed a mixed-effects model in which fixed effects nPE magnitude, nPE valence, condition (human or computer) and their respective interaction terms and a random effect for unit predicted group activity of SNr putative DA units during the 1000 ms after split reveal for all SNr data pooled data together (S9 Table). This model revealed that mean firing rates of SNr putative DA units were significantly predicted not only by nPE valence (nPE valence, *\*\*p* = 0.0057, **β** ± SEM = 0.20 ± 7.3e-02, 95% CI [0.06, 0.34]), but also by the interaction between nPE valence and condition (nPE valence x condition, *\*p* = 0.036, **β** ± SEM = -0.22 ± 0.11, 95% CI [-0.43, - 0.01]), such that the encoding of nPE valence by firing rate was significantly stronger in the human condition than in the computer condition.

Finally, to account for the potential role of offer size in the observed neural response to nPE valence, we constructed a linear mixed-effects model predicting SNr putative DA unit activity at split reveal in the human condition by fixed effects offer size and nPE valence with a random effect for unit. This analysis revealed that nPE valence remained a significant predictor of firing rate at split reveal even after controlling for offer size (nPE valence, *\*p* = 0.022, **β** ± SEM = 0.12 ± 5.1e-02, 95% CI [0.02, 0.21]), while offer size was not a significant predictor (offer, *p* = 0.45, **β** ± SEM = -3.6.e-03 ± 4.7e-03, 95% CI [-0.01, 0.01]). We repeated this analysis, instead treating offer as a categorical variable. Again, nPE valence was the only significant predictor of firing rate (nPE valence, *\*p* = 0.030, **β** ± SEM = 0.11 ± 5.1e-02, 95% CI [0.01, 0.21]), with no significant effect of offer. These findings suggest that the observed firing activity of these putative DA neurons is more consistent with encoding of whether an offer is better or worse than the participant’s internal norm expectation than with offer magnitude alone.

Altogether, these findings indicate that putative SNr DA units encode nPE valence, with greater encoding observed in the human condition, providing a unique neural mechanism by which SNr neurons track social information.

### Putative neurons in the GPi do not encode social prediction error valence

It is possible that the nPE encoding effect observed in SNr putative single-unit activity reflects widespread activation across basal ganglia (BG) nuclei. To examine the anatomical specificity of this effect, we carried out a similar experiment in another cohort of neurosurgery participants who underwent DBS surgery in a different BG nucleus, the GPi (5 men, n = 3 PD and n = 2 dystonia patients, aged 53 ± 8.5 years; **S9A Figure**). From these 5 participants, we collected 5 sessions of UG with human block only and 2 sessions of UG with both human and computer blocks and mood ratings. In this group, rejection rate decreased with increasing offer size with no significant differences between conditions (**S9B Figure**). A logistic regression model predicting binary choices by offer size found a significant effect (offer, \*\*\**p* = 1.9e- 14, **β** ± SEM = 1.3 ± 0.17), indicating a higher likelihood of accepting higher offers than lower offers. To facilitate comparison with the SNr recording group, we estimated nPEs for these sessions using the winning model from the SNr cohort, namely the RW variable initial norm model with fixed alpha.

We identified 19 putative units from 7 recording sessions (mean ± SEM clusters per recording: 2.7 ± 0.29). Waveform widths and average firing rates for all 19 putative GPi neurons are listed in **S10 Table**. The waveforms for all 19 putative GPi units and the mean waveform for these units are depicted in **S9C Figure.** We smoothed spike trains into continuous firing rates according to the protocol implemented in the SNr unit analyses. We then related GPi putative unit activity during the 1000 ms after split reveal to nPE valence, nPE magnitude, and their interaction by a mixed-effects regression identical to the regression used to predict SNr putative DA unit activity at split reveal in the human condition. This model was not constructed in the computer condition given that only two players completed UG with a computer block. Note also that 3 of 19 putative units were recorded during a behavioral session in which all offers were accepted; so, behavioral modeling was not performed on this session, resulting in 16 total putative GPi units for analysis in relation to nPE valence, magnitude, and their interaction. This regression revealed no significant effects for any of these variables or their interactions (**Table 3**).

**Table 3.** Summary of mixed-effects model predicting all globus pallidus internus (GPi) putative neuron activity at split reveal.

| Fixed effect | Estimate | Std. error | t-value | p-value |
| --- | --- | --- | --- | --- |
| **Human** |  |  |  |  |
| Intercept | 0.09446 | 0.04858 | 1.944 | 0.0545 |
| nPE magnitude | -0.01331 | 0.01224 | -1.087 | 0.2775 |
| nPE valence | 0.17970 | 0.15304 | 1.174 | 0.2409 |
| nPE magnitude x nPE valence | -0.18235 | 0.09359 | -1.948 | 0.0519 |

Summary of mixed-effects model in which fixed effects norm prediction error (nPE) magnitude, nPE valence, and their respective interaction terms predicted group activity of all GPi units during the 1000 ms after split reveal in the human condition. Std. Error = standard error. $n = 16$ putative neurons in the human condition.

In order to facilitate a comparison between the results of the GPi neurons and those of the SNr neurons that is less biased by waveform and firing rate differences between populations, 8 of these 16 GPI putative units were selected for a repeat analysis in which firing rate and waveform length criteria were matched to those of the putative DA SNr units. We again related GPi putative unit activity during the 1000 ms after split reveal to nPE valence, nPE magnitude, and their interaction terms by mixed-effects regression in the human condition only. This regression revealed no significant effects for any of these variables or their interactions (**Table 4**). (For visualization of mean firing rates at split reveal by nPE valence in the human condition, see **S6D Figure**.) Lack of a statistically significant effect in this population could certainly be related to the limited sample size of recorded units in this region. So, with this limitation in mind, our results suggest that while our sample of putative DA neurons in the SNr demonstrate encoding of nPE valence, this pattern was not observed with significance in our sample of putative units from the GPi, even when matched by waveform and firing rate criteria.

**Table 4.** Summary of mixed-effects model predicting waveform- and firing rate-matched globus pallidus internus (GPi) putative neuron activity at split reveal.

| Fixed effect | Estimate | Std. error | t-value | p-value |
| --- | --- | --- | --- | --- |
| **Human** |  |  |  |  |
| Intercept | 0.088536 | 0.080590 | 1.099 | 0.2771 |
| nPE magnitude | -0.004832 | 0.020630 | -0.234 | 0.8150 |
| nPE valence | 0.478699 | 0.268627 | 1.782 | 0.0761 |
| nPE magnitude x nPE valence | -0.276125 | 0.164248 | -1.681 | 0.0941 |

Summary of mixed-effects model in which fixed effects norm prediction error (nPE) magnitude, nPE valence, and their respective interaction terms predicted group activity of GPi units (matched in waveform and firing rate criteria to those of the SNr putative DA unit group) during the 1000 ms after split reveal in the human condition. Std. Error = standard error. $n = 8$ putative neurons in the human condition.

### Putative DA neurons in the SNr do not encode mood signals

Finally, given that PEs have previously been demonstrated to impact momentary mood,^16,41^ we examined whether the activity of putative DA SNr neurons also tracked subjective mood. We observed that putative DA unit firing rates during the 1000 ms before mood ratings were not significantly correlated with z-scored mood ratings (Pearson’s product-moment correlation, *p* = 0.16, r = -0.09, 95% CI -0.21 - 0.03) (**S10 Figure**). A mixed-effects model predicting DA unit group activity during this period by mood rating with a random effect for unit revealed no significant effect for mood rating (mood rating, *p* = 0.16, **β** ± SEM = - 0.027 ± 0.019), consistent with the results of the correlation test. These findings suggest that, while putative DA SNr neurons respond to nPE valence, they do not encode mood ratings.

## DISCUSSION

The current study examined how single neurons in the human SNr encoded social learning and mood signals. Behaviorally, we found that participants frequently rejected non-zero offers, consistent with previous work demonstrating that rejection behavior in UG is driven not by objective utility, but rather by subjective utility, which reflects how valuable an offer is to a specific individual, given his or her norm expectations and personal sensitivity to unfair inequality.^14,15^ These behaviors persisted even during gameplay with computer avatar partners, indicating some degree of social thinking in less social contexts, as indicated by the outperformance of social norm models in explaining participant choices when compared to non-social reward learning. Social norm modeling revealed that participants had greater expectations for fairness from humans versus computers,^37^ and nPEs influenced their momentary mood.^16,41^ Neurally, substantia nigra neurons tracked the valence of experienced nPEs, with greater encoding in the human avatar condition; these neurons did not encode momentary mood. Collectively, these findings pinpoint a key role of the substantia nigra in encoding social learning signals.

First, our results demonstrate the utility of computational modeling in revealing how similar observed behaviors may arise from distinct underlying mental processes. Although rejection rates were similar across human and computer avatar conditions, our model demonstrates that these choices were informed by latent variables including initial norm expectations, sensitivity to unfairness, choice variability, and norm adaptation. Specifically, higher initial and session-wide internal norms were observed during gameplay with human versus computer avatar partners, indicating that expectations for fairness, which in turn impact choice behavior, were indeed contingent on counterpart identity. As a result, this modeling approach, strengthened by strong parameter recovery and posterior predictive checks, elucidates meaningful latent processes influencing choice behavior that would not have been apparent based on raw rejection rates, alone.

Moreover, our results highlight an important role of SNr neurons in encoding valence across the domains of social and non-social learning. Previous work has identified roles for the SN in reward processing and reinforcement learning,^17–23,28^ with DA activity implicated in prediction error encoding in nonsocial contexts.^29–31^ One such study conducted within neurosurgical patients demonstrated that a population of SN putative dopaminergic neurons exhibits higher firing rates in response to unexpected gains versus losses, without evidence of a statistically significant relationship between firing rate and magnitude, similar to encoding of nPE valence in social contexts observed in our study.^22^ Additionally, midbrain DA neurons in non-human primates have been observed to respond specifically to positive error signals in nonsocial contexts.^18^ Our findings expand upon this work by providing evidence in awake humans that these reward-associated single SNr neurons also show heightened activity in response to positive versus negative nPEs during gameplay with human avatars, demonstrating that these putative DA units encode prediction error valence specifically in the social domain. The observed encoding of valence in the spiking rate of putative DA SNr neurons is likely to primarily impact DA concentration in the striatum through ascending axonal projections.^43^ This activity may also impact nigral activity through local dendritic release of DA, consistent with recent previous work identifying heightened DA concentration in the SNr during interactions with human versus computer avatars using human voltammetry.^32^ Further, while the putative dopaminergic SNr neurons recorded in our study respond specifically to nPE valence, our findings do not negate that other neurons may encode magnitude.

Consistent with previous work relating prediction errors to mood,^16,41^ our findings demonstrate that, while nPEs influence momentary mood states, and the valence of these nPEs is encoded in SNr neural activity, changes in mood are likely reflected in neural activity elsewhere in the human brain.^44^ Our findings expand upon recent human studies relating SN structural and functional abnormalities to emotional disturbances and symptoms,^45,46^ and intracranial signals to social valuation signals,^47^ by contributing evidence from directly recorded single SNr neurons in awake humans during emotionally upsetting social interactions. We thus contribute to the SN literature by outlining a role for these neurons in connecting learning signals with mood in social contexts.

Notably, our analyses did not reveal significant neural tracking of nPE signals by the GPi, a neighboring basal ganglia nucleus. This finding is consistent with work demonstrating very sparse responses of single units to reward this region.^48^ .

Our findings should be interpreted with consideration of the following limitations. The study participants are largely male and of advanced age, which is characteristic of those who are most likely to undergo DBS surgery for PD.^49^ Our study therefore may not account for sex- or age- related differences in social norm processing. These participants also all have PD, which is characterized by behavioral rigidity,^50^ mental inflexibility,^51^ and reduced ability to switch cognitive strategies when those strategies are no longer favorable.^52^ PD pathology additionally impacts our targeted brain regions, causing hyperactivity in SNr and GPi neurons due to reduced dopaminergic input.^53,54^ These patients participate in our study while off dopaminergic medications, which facilitates examination of the role of surviving putative dopaminergic SN neurons in social norm learning without the inference of medications that may alter learning and decision making; however, the dopaminergic signaling in these patients occurs within an inherently dopamine depleted state, which means that the results of this study, while novel, may not be generalizable to the dopaminergic circuitry of healthy individuals. To better ascertain how the behavioral and neural changes induced by PD pathology may influence the results obtained from our study, it would be most suitable to compare behavioral and neural data obtained from individuals with and without PD. While behavioral results can be compared to those obtained from other participants of studies using UG, single-unit neural data is not as readily available for comparison, given the highly invasive nature of these recordings. For similar reasons, this study is limited in its sample size of participants. Additionally, in an effort to use all relevant data available from this limited sample of participants, this study incorporates data collected from those who played both the human and computer conditions of the task as well as from those who played only the human condition, which creates imbalance in condition exposure. This study is also limited in its ability to define precisely how recorded single neuron activity in the SN translates into dopamine levels in the striatum; such a delineation could be made possible with single unit SN recording and simultaneous voltammetric recording in the striatum.^43^ Finally, this study does not demonstrate a causal role for these neurons in social norm processing; future work using stimulation or other perturbation techniques could address causality directly.

Taken together, our work suggests a neural basis for the human ability to recognize errors in social prediction, supporting a role for the SN not only in classically studied reward learning but also in the learning of normative social behaviors.

## MATERIALS AND METHODS

### Ethics and participants

This study was approved by the Institutional Review Board at the Icahn School of Medicine at Mount Sinai. No adverse events occurred as a result of this study. 24 neurosurgery patients (21 men, 3 women, mean age ± SEM = 62 ± 2.4 years; **S1 Table**) participated in this study. After consenting to deep brain stimulation (DBS) surgery, patients deemed suitable for research participation by our lead neurosurgeon were then given the option to learn about participation in this study. If interested in hearing more, patients were given information about the study, its impact on the procedure, and any potential risks. They were also given a brief introduction to the behavioral task, including instructions and two trial examples. Patients were informed that they would not receive monetary compensation for their participation. Rather, they were instructed to imagine splitting $20 iteratively with human or computer avatar partners. All information was provided both verbally and in written form with ample time for questions and clarifications before receiving written consent.

### DBS surgery and single-unit recording

DBS device implantation occurred over three surgeries, including left hemisphere lead implantation, right hemisphere lead implantation, and battery implantation. These surgeries were approximately 2-4 weeks apart. Participants were given the opportunity to participate in research during either or both lead implantation surgeries. At the time of surgery, participants were off their medications for at least 9 hours, as mandated by standard clinical protocol.

At the point in DBS surgery when patients were woken up for clinical lead targeting and testing (which typically occurred approximately 10 minutes after lead implantation), the clinical procedure was paused briefly for single unit recording and task participation. Our lead neurosurgeon positioned the tungsten microelectrode into the recording target using power-assisted microdrive and tractography. After positioning the microelectrode within recording target regions as identified by their unique signatures of neural firing, the neurosurgeon further manipulated electrode position until confirming the presence of stable spike activity. Neural data was recorded using an Alpha Omega electrophysiological recording system. For patients who were undergoing DBS surgery in the subthalamic nucleus, single-unit recordings were collected from the substantia nigra pars reticulata (SNr; **Figure 1A**). For patients undergoing DBS surgery in the globus pallidus internus (GPi), single-unit recordings were also collected from the GPi **(S9A Figure**). After recordings were collected, the clinical stimulator was repositioned to the clinical target region for testing and final implantation.

The SNr cohort included 19 participants with Parkinson’s disease (16 men, 3 women, aged 65 ± 1.93 years). The GPi cohort included 5 participants with Parkinson’s disease or dystonia (5 men, 0 women, aged 53 ± 8.5 years). During single-unit recording, participants played a behavioral task visualized on a computer monitor positioned within eyesight. Participants submitted their answers with a game controller box. In total, we collected 27 sessions from the SNr cohort and 7 sessions from the GPi cohort.

### Ultimatum game

During each single-unit recording session, participants played an Ultimatum Game (UG), which is used to assess social fairness norms^37,38^ and norm adaptation^14^ (**Figure 1B**). Participants played one of two versions of UG: human-only (n = 15 sessions in 10 patients from the SNr cohort; n = 5 sessions in 3 patients from the GPi cohort) and full version with both human and computer conditions (n = 12 sessions in 9 patients from the SNr cohort; n = 2 sessions in 2 patients from the GPi cohort).

In the human-only version, participants are repeatedly presented with proposed splits of $20 by “human” avatar partners, which they could accept or reject. If a participant accepted, each party received the proposed amounts. If a participant rejected, each party received $0. At the start of each trial, participants were first presented with an avatar (1.5-2.5 s). Next, participants were presented with the proposed split of $20 (“split reveal”; 1 sec). Choice buttons (accept/reject) then appeared below the proposed split, prompting participants to make their selection (self-paced). After making their selection, the selected choice remained highlighted for 1 second before the screen changed to reveal the resulting dollar amounts received by each party (“split result”; 1.5 s). All offers were $9 or less, and offers were pseudo-randomized with a normal distribution between $1 and $9. Choices did not impact future offers. Players were not privy to the offer structure in advance. At each trial, a new offer was proposed by a new human avatar. Human avatars varied by name, gender, and physical features in efforts to portray racial and cultural diversity. Patients completed 30 trials, which took approximately 5 minutes to complete.

In the full version with both human and computer conditions, participants played 30 trials of UG with human avatar partners, as well as an additional block of 30 trials with “computer” partners. Block order was counterbalanced such that 5 participants played the human block before the computer block, and 7 participants played the computer block before the human block. Additionally, participants completed a self-reported mood rating (self-paced) after a random 33% of choice trials. To complete this mood rating, participants selected their present mood on a scale from an unhappy face (coded as 0) to a happy face (coded as 100). Two minor screen presentation details differed slightly from those in UG: split reveal appeared simultaneously with choice buttons, and split result lasted for 1 s instead of 1.5 s. Altogether, this version of UG took approximately 10 minutes to complete.

### Exclusion criteria

To protect data quality from compromise due to inattention or drowsiness, all sessions in which behavior indicated a failure to follow task instructions were excluded from behavioral analysis. These included sessions with flat behavior, defined as sessions in which less than 2 offers are accepted or rejected in the entirety of a human or computer block. This measure effectively excluded sessions in which participants accepted all offers, which prevents the choice variability required for model-fitting. Excluded sessions also included those in which logistic fits of binary choice versus offer size have non-positive slopes, meaning that probability of accepting an offer does not increase with increasing offer size. If mood ratings were flat, mood ratings were excluded from analysis, but choice behavior was still included. Behavioral sessions with flat mood ratings are those in which minimum and maximum ratings do not differ by at least 10. In the SNr cohort, 3 UG and 3 UG-bot sessions were excluded from behavioral analyses due to flat choices. An additional 2 UG-bot sessions were excluded from mood rating analyses due to flat ratings. In the GPi cohort, 1 UG session was excluded from behavioral analyses due to flat choices, and 1 UG-bot session was excluded from mood rating analyses.

### Computational modeling of behavior

All data cleaning, behavioral analyses, and computational models were implemented in MATLAB and RStudio. Model-free analyses included: overall rejection rate by block type (human/computer), rejection rate vs. offer size, overall reaction time by block type, overall mood rating by block type, and mood rating vs. choice (accept/reject).

Next, models previously implemented^14,15^ to model utility, decision probabilities, and norm adaptation were fitted to choice data. First, we represented the utility of the exchange on each trial using the Fehr-Schmidt (FS) equation for inequality aversion utility:^15^

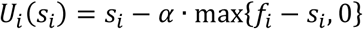

Here, the utility (*U_i_*) of an offer (*s_i_*) equals the difference between *s_i_* and the product of the difference between the participant’s internal norm (*f_i_*) and *s_i_*and the participant’s sensitivity to disadvantageous inequality (α ∈ [0,1], “alpha”). Alpha reflects the participant’s unwillingness to accept an offer below the internal norm at that trial. Higher values of α represent greater aversion to unfairness. Sensitivity to advantageous inequality (“guilt”) is unnecessary in this paradigm, since all offers are $9 or less.

We then modeled the probability of accepting each offer using a softmax function:^14^

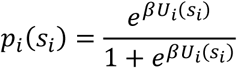

This function considers the utility of each offer and an inverse temperature (*β ∈* [0,1]) parameter, which reflects the variability of the participant’s choices. Low values of *β* represent more variable choices.

In FS models, the norm is considered a static, non-changing metric.^13–15^ To model internal norm adaptation, we compared two classes norm adaptation models for best fit to this cohort’s behavior: Rescorla-Wagner (RW) models, which are derived from temporal difference learning,^14,39,40^ and Bayesian observer (Bayes) models.^13,14^ Both classes of norm adaptation models assume that the internal norm evolves over time; they differ, however, in their norm updating rule.

The RW models are a reinforcement learning based approach in which an agent learns to take actions that maximize reward within an uncertain environment.^14,39,40^ In the RW models, the norm is updated as follows:

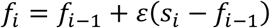

Here, the internal norm (*f_i_*) at trial *i* equals the norm at the previous trial (*f_i-1_*) plus the product of norm adaptation rate (*ε* ∈ [0,1]) and the difference between an offer (*s_i_*) and the internal norm at the previous trial (*f_i-1_*), or the nPE. *ε* reflects the magnitude of the influence of the offer (*s_i_*) on the evolving internal norm (*f_i_*).

The Bayesian models, on the other hand, are derived from a statistical approach that estimates a probability distribution over model parameters based on observed data.^13,14^ We assume that each participant models the offers as originating from a normal distribution characterized by unknown mean and variance. The participant has an initial prior of the mean (μ) and variance (σ^2^), and this prior is updated with every new offer. Given μ and σ^2^, the offers (X) are defined as: *X |* μ,σ^2^ ∼ *N* (μ,σ^2^). After receiving the first offer (*s_i_*), and assuming the prior *p*(μ,σ^2^), the posterior is defined as:^14^

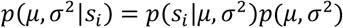

The conjugate prior is defined as:^14^

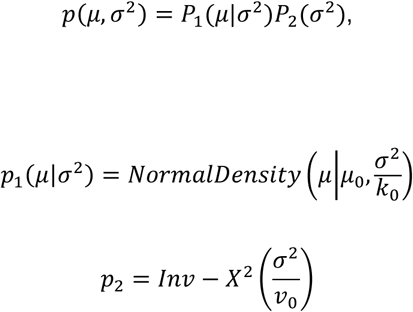

Next, after all subsequent offers, the posteriors are defined iteratively by:^14^

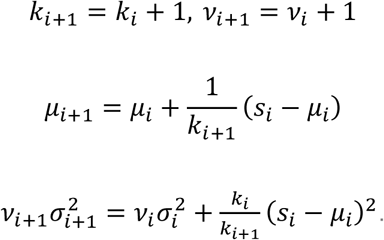

Finally, on trial *i* + 1 the updated internal norm is defined by:^14^

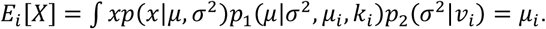

Within each class of model, individual models differed by their assumptions about the initial internal norm; models either assumed a fixed initial norm expectation equal to 10 (corresponding to an equal split of $20; labeled “fixed f0”) or a variable initial norm expectation that was individually fit to each subject’s behavior (labeled “variable f0”). The variable initial norm models also differed in whether alpha was individually fit to each subject’s behavior or fixed across the patient sample to the group’s mean alpha (labeled “fixed alpha”).

All models were fitted to each subject’s behavioral sessions (with human and computer blocks treated separately) and compared for best fit as determined by minimum total Bayesian information criterion (BIC) score across the group. Total BIC is the group-level sum of BIC scores for each subject for each model. The FS: fixed alpha model won, followed by the RW: fixed alpha model (**Figure 2D**).

Next, to assess whether behavior is driven by a classic RPE, we compared all 9 models to a Q-learning approach.^16^ A Q-learning algorithm learns an optimal action-value function within an environment without any prior model of the dynamics of that environment. It does so by updating Q-values, which are estimates of the value of taking an action in a given state based on received reward and the estimated value of the next state. This method adjusts Q-values incrementally as driven by RPEs. RPE-directed updates are made at a learning rate between 0 and 1 after the reward is received and before the next offer is proposed. The initial Q-values are as follows:

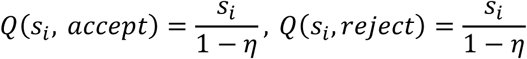

These Q-values give the initial value of a state-action pair in which an offer *s_i_* is accepted or rejected. Eta (η) represents the temporal discounting factor, which determines how much future rewards influence the current value; here, η is 0.8.^16^

Next, for trial i > 1, based on the previous choice (i.e. accept or reject) at trial i-1, the Q-learning update rule if an offer is accepted is the following:^16^

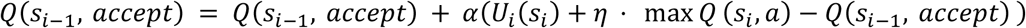

Here, alpha (α) is the learning rate which determines how much the prediction error updates the old Q- value. α is between 0 and 1; 0 reflects no update (i.e. no learning), while 1 reflects a complete update (i.e. fast learning).

Alternatively, the Q-learning update rule if an offer is rejected is the following:^16^

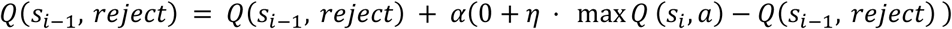

The value difference is:^16^

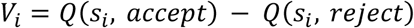

Finally, the probability of accepting and offer is then modeled by:^16^

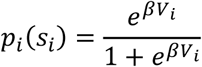

while an nPE describes how much higher or lower an offer is above or below an individual’s model- based norm for fairness, an RPE, on the other hand, represents how much higher or lower an outcome was versus an expectation based on previous events.

We compared the BIC scores of all 9 models to the BIC score of the Q-learning approach and found that the FS: fixed α model again produced the best fit, followed by the RW: fixed α model, with the Q- learning approach performing worse than all 9 other models (**S2A Figure**). These results were consistent across both across and between human and computer conditions (**S2B-C Figure**), suggesting that behavior in our task is motivated not by classic Q-learning, but rather by statis social preferences as in the FS: fixed α model, or by model-driven incorporation of new information into norm expectations over time as driven by nPEs, as in the RW: fixed α model.

Next, to facilitate better judgment of the strength of evidence for each of the two best-fitting models, we assessed ΔBIC, expected model frequency, and protected exceedance probabilities (S3 Table). These analyses demonstrated that the FS: fixed α model had the lowest BIC with an expected model frequency of 0.435 and a protected exceedance probability of 0.932. The RW: fixed α model had a ΔBIC of 13.707, expected model frequency of 0.174, and protected exceedance probability of 0.043. This means that 43.5% of the population is expected to be best represented by the FS: fixed alpha model, while 17.4% of the population is expected to be best represented by the RW: fixed alpha population. So, while there is greater evidence in support of the FS: fixed α model overall, there is also evidence of some heterogeneity in best fit between the two candidate models, consistent with the fact that mean BIC scores were comparable at 10.45 and 10.93 for the FS: fixed α and RW: fixed α models, respectively.

We then continued our assessment of these two models by exploring the degree to which each model was identifiable via model recovery (S3 Figure; **S4 Table)**. First, we computed the empirical mean and standard deviation for each parameter of each of the 10 models, constructed normal distributions centered on those estimates for each parameter within each model, and drew new parameter values from those distributions, via the same parameter sampling procedure used in our parameter recovery. For each generating model, we simulated behavioral datasets for 29 synthetic subjects using these sampled parameters, corresponding with the 29 empirical behavioral datasets that underwent model fitting in our study. We then refit all 10 models to these simulated datasets. This process was repeated for 200 independent repetitions, with newly sampled parameter values and simulated datasets generated for all 29 synthetic subjects in each repetition. Afterward, we constructed both a 10x10 recovery matrix and a 4x4 class-level recovery matrix (RW models, Bayes models, FS models, and the Q-learning model) to further address the question of whether these model classes represent distinguishable computational processes.

At the individual model level, the highest recovery probabilities were demonstrated by the Q-learning (50.79%), Bayes: fixed alpha (45.88%), FS: fixed f0 (45.83%), and RW: fixed alpha (45.47%) models (see new S4 Table and S3 Figure added to the manuscript and copied below). The FS: fixed alpha model demonstrated weaker recovery, with a recovery probability of 37.62%. Finally, the RW: variable f0, RW: fixed f0, Bayes: variable f0, Bayes: fixed f0, and FS: variable f0 were essentially unrecoverable (recovery probabilities 1.28%, 9.72%, 0.41%, 9.33%, and 0.71%, respectively). These results demonstrate that the variable f0 models tend to be less identifiable, while the fixed α models tend to be more identifiable. Further examination of the recovery matrix revealed that confusion most often occurred between related models within the same model class.

To evaluate whether models were more distinguishable at the class level, we constructed a class-level recovery matrix by averaging the recovery probabilities across models within each class. At the class level, recovery probabilities for RW models, Bayes models, FS models, and the Q-learning model were 40.57%, 47.67%, 81.82%, and 50.79%, respectively (see new S5 Table and S4 Figure added to the manuscript and copied below). Most confusion occurred between the RW and Bayes families, consistent with the fact that both classes implement model-based norm-adaptation mechanisms. Overall, these analyses demonstrate that, while related individual models are sometimes confused with one another within shared model classes, these higher-level model classes are more reliably recovered, indicating that the behavioral predictions generated by these computational model classes are fairly distinguishable. Moreover, out of the two best-fitting models, the RW: fixed α model demonstrated the best identifiability on model recovery.

### Parameter recovery

Next, to test for recoverability of the parameters generated by the RW: fixed *α* model, 29 behavioral sessions were artificially simulated based on normal distributions of the parameter estimates obtained from the 29 empirical behavioral sessions. The simulated behavior was then subjected to the same modeling process as the original empirical behavior. Resulting model parameters obtained from the simulated fits were then compared by correlation to the original empirical parameter estimates. Higher Pearson correlation coefficients indicated better parameter recoverability. All model parameters were significantly recovered (**S5 Figure**; temperature: r = 0.51, \*\**p* = 0.0049; norm adaptation rate: r = 0.86, \*\*\**p* = 1.8e-09; initial norm: r = 0.94, \*\*\**p* = 6.9e-14). We therefore concluded that this norm adaptation model can be applied with adequate replicability.

### Posterior predictive checks

We next examined whether the RW: fixed *α* model reproduced the choice patterns observed in our study. To do so, we simulated 29 behavioral datasets given the parameter estimates obtained from empirical model fitting with the RW: fixed *α* model, using the exact offer vectors presented in each session. The RW: fixed *α* model reproduced overall rejection rates in both the human (mean ± SEM: observed: 0.662 ± 0.042; simulated: 0.652 ± 0.045) and computer (observed: 0.542 ± 0.050; simulated: 0.500 ± 0.065) avatar conditions(**S6A Figure).** The RW: fixed *α* model also recapitulated the negative impact of offer size on rejection rate (**S6B Figure)**. Finally, to assess whether the RW: fixed *α* model recapitulates behavior at the individual subject level, individual rejection rates for observed versus simulated sessions were also compared, revealing a strong linear correlation (**S6C Figure**; Pearson’s product-moment correlation, \*\*\**p* = 4.55e-12, r = 0.914, 95% CI 0.823 - 0.959).

### Comparison of parameters by condition

For the RW: fixed alpha model, parameter estimates for inverse temperature, norm adaptation rate, and initial norm were compared by mixed-effects models predicting each parameter by condition (human or computer avatar) with random effects for participant and for session within participant (**Table 1**). A mixed-effects model predicting temperature by condition with random effects for participant and for session within participant found no significant effect of condition (condition, *p* = 0.34, **β** ± SEM = 0.80 ± 0.83; OR = 2.24, 95% CI [0.41, 12.17]). A mixed-effects model predicting initial internal norm by condition with random effects for participant and for session within participant found a significant effect for condition such that initial norms were higher during gameplay with human versus computer avatar partners (condition, \**p* = 0.025, **β** ± SEM = -4.76 ± 2.00; OR = 0.0085, 95% CI [0.00014, 0.52]). Finally, a mixed-effects model predicting norm adaptation rate by condition with random effects for participant and for session within participant found no significant effect of condition (condition, *p* = 0.083, **β** ± SEM = 0.053 ± 0.030; OR = 1.05, 95% CI [0.99, 1.12]). So, while temperature and norm adaptation rate did not differ by condition, initial norms did indeed differ by condition, such that norms for fairness were higher during gameplay with human avatar partners.

### Analysis of single-unit electrophysiological data

Neural data was digitized at 44,000 Hz and filtered at 300-9000 Hz to select only high frequency activity. We identified clusters from filtered single-unit recordings in MATLAB using the OSORT offline spike sorting algorithm developed by Rutishauser, et al.^42^ Spikes were detected only when the neural signal exceeded the threshold of the standard deviation of the average signal amplitude multiplied by five. As each new spike was detected, that spike was compared by multidimensional distance to clusters of similar, previously detected spikes. If the distance was smaller than the clustering threshold (the average standard deviation squared of the filtered signal along a sliding window multiplied by the number of data points in a waveform) then it was added to that cluster; if the distance was larger than the clustering threshold, then it was categorized as a new cluster.

We then manually examined each cluster returned by the sorting algorithm, eliminating artifacts and merging clusters that were falsely split. A cluster was considered an artifact if >5% of all spikes originating from that cluster had an interspike interval (ISI) of less than 3 ms, corresponding to the length of the refractory period.^42^ False splits were identified by examining the projection tests generated by OSORT for each cluster. These projection tests provide visualizations of the multidimensional distance between each possible pair of clusters.

We categorized clusters characterized by an ISI <5% as putative single units.^42^ We generated raster plots and peristimulus time histograms for each putative unit at split reveal, the point in UG at which the offer is presented. We then smoothed spike trains for putative units into continuous waveforms using a sliding, symmetric Gaussian kernel (width = 300 ms) centered at each time point (see **S8 Figure** for one example unit). Smoothing was zero-phase and did not introduce temporal shifts in the timing of neural responses. We computed average firing rate for each putative unit and characterized average waveforms by number of peaks, waveform width, polarity, and distance between peaks and troughs. In the SNr, we identified 69 putative neuronal clusters from 27 SNr microelectrode recordings (mean ± SEM clusters per recording: 2.6 ± 0.11). We labeled units with firing rates less than or equal to 15 Hz and waveform widths greater than 0.8 ms with biphasic shape as putative dopaminergic (DA) units.^27^ If a unit’s bipolar waveform had a negative deflection followed by a positive deflection (rather than the reverse polarity), then this unit was considered a putative DA unit likely reversed in orientation relative to the position of the electrode. In total, 40 of 69 putative units were labeled as putative DA units (mean ± SEM putative DA units per recording: 1.5 ± 0.18) (**Figure 3A**). The waveforms for all 40 putative DA units and the mean waveform for these units are depicted in **Figure 3B**. Strictly for visualization purposes in **Figure 3B**, waveforms for putative DA units with reversed polarity were reflected over the x-axis (i.e. microvolts multiplied by –1 at each time point) to correct for flipped orientation relative to the electrode tip. Continuous firing rates for each unit were z-scored across the entire recording for each unit, using the mean and standard deviation of the unit’s activity across the full session. These z-scored continuous firing rates were then extracted around events of interest (i.e. split reveal, mood rating, etc) for subsequent population-based analyses.

To test whether the activity of these putative DA neurons might encode properties of nPEs, we first compared the firing rates of these units during the 1000 ms after split reveal during trials in which nPEs were positive versus those in which they were negative. Group-level putative DA unit activity appeared to be higher during positive nPE trials versus negative nPE trials in the human condition (for probability density functions see **Figure 3E**); this pattern was attenuated in the computer condition (for probability density functions see **Figure 3F**). To test the statistical significance of this observation, and to also test whether these units encode nPE magnitude, we implemented a mixed-effects model to predict the activity of these units during the 1000 ms after split reveal by nPE valence (1 = negative, 2 = positive), magnitude, and their interactions with a random effect for unit in each condition. Valence was treated as a factor. All mixed-effects models were implemented using the “lme4” and “lmerTest” packages in RStudio. The mixed-model for the human condition resulted in a significant effect for valence (nPE valence, \*\**p* = 0.008, **β** ± SEM = 0.19 ± 7.3e-02, 95% CI [0.05, 0.34]; **Figure 3G**; see **Table 2** for other estimates). Valence was not a significant predictor of firing rate in the computer condition (PE valence, *p* = 0.79, **β** ± SEM = -0.02 ± 7.7e- 02, 95% CI [-0.17, 0.13]; **Figure 3H**; see **Table 2** for other estimates).

To formally test how the nPE valence effect differs across conditions, we constructed a mixed-effects model in which fixed effects nPE magnitude, nPE valence, condition (human or computer) and their respective interaction terms and a random effect for unit predicted group activity of SNr putative DA units during the 1000 ms after split reveal for all SNr data pooled data together (S9 Table). This model revealed that mean firing rates of SNr putative DA units were significantly predicted not only by nPE valence (nPE valence, *\*\*p* = 0.0057, **β** ± SEM = 0.20 ± 7.3e-02, 95% CI [0.06, 0.34]), but also by the interaction between nPE valence and condition (nPE valence x condition, *\*p* = 0.036, **β** ± SEM = -0.22 ± 0.11, 95% CI [-0.43, - 0.01]), such that the encoding of nPE valence by firing rate was significantly stronger in the human condition than in the computer condition. Together, these results demonstrate that putative SNr DA units may be sensitive to nPE valence, especially in more social contexts.

To account for the potential role of offer size in the observed neural response to nPE valence, we constructed a linear mixed-effects model predicting SNr putative DA unit activity at split reveal in the human condition by fixed effects offer size and nPE valence with a random effect for unit. This analysis revealed that nPE valence remained a significant predictor of firing rate at split reveal even after controlling for offer size (nPE valence, *\*p* = 0.022, **β** ± SEM = 0.12 ± 5.1e-02, 95% CI [0.02, 0.21]), while offer size was not a significant predictor (offer, *p* = 0.45, **β** ± SEM = -3.6.e-03 ± 4.7e-03, 95% CI [-0.01, 0.01]). We repeated this analysis, instead treating offer as a categorical variable. Again, nPE valence was the only significant predictor of firing rate (nPE valence, *\*p* = 0.030, **β** ± SEM = 0.11 ± 5.1e-02, 95% CI [0.01, 0.21]), with no significant effect of offer. These findings suggest that the observed firing activity of these putative DA neurons is more consistent with encoding of whether an offer is better or worse than the participant’s internal norm expectation than with offer magnitude alone.

Finally, we examined how the activity of these putative DA units relates to momentary mood ratings. We first implemented a Pearson’s product-moment correlation test between z-scored mood ratings and putative DA unit firing rates during the 1000 ms before mood rating selection, finding that firing rates and z-scored mood ratings were not significantly correlated (*p* = 0.16, r = -0.09, 95% CI -0.21 - 0.03) (**S10 Figure**). We next implemented a mixed-effects model to predict putative DA unit group activity during this epoch by z-scored mood rating with a random effect for unit, which confirmed no significant impact of mood rating on neural activity (mood rating, *p* = 0.16, **β** ± SEM = -0.027 ± 0.019). These results suggest that the activity of these putative DA units responds uniquely to the valence of nPEs but not to mood ratings.

## RESOURCE AVAILABILITY

### Lead contact

Requests for further information and resources should be directed to and will be fulfilled by the lead contact, Arianna Davis.

### Materials availability

This study did not generate any unique reagents.

### Data and code availability

- De-identified behavioral and neural recording data have been deposited at figshare and are publicly available as of the date of publication at https://doi.org/10.6084/m9.figshare.29082671.v2.
- All original code has been deposited at figshare and is publicly available at https://doi.org/10.6084/m9.figshare.29082671.v2 as of the date of publication.
- Any additional information required to reanalyze the data reported in this paper is available from the lead contact upon request.

## ACKNOWLEDGMENTS

We thank the members of the Gu and Saez laboratories for advice and feedback during manuscript review and editing. No AI-assisted technologies were used in preparation of the manuscript. The data in this paper were used to complete a dissertation as partial fulfillment of the requirements for a PhD degree at the Graduate School of Biomedical Sciences at Mount Sinai.

## AUTHOR CONTRIBUTIONS

Conceptualization: XG, IS Data curation: AND

Formal analysis: AND, SEQ, XG, IS Funding acquisition: XG, PRM, AND, IS

Investigation: AND, MH, NM, ZI, AK, BHK, OP

Methodology: AND, SN, XG, SEQ, ZI, BHK, PRM, IS, QXF

Project administration: AND, XG, IS, MH

Resources: XG, IS, BHK, NM, PRM

Software: QXF

Supervision: XG, IS, HSM

Visualization: AND, XG, IS

Writing – original draft: AND, XG, IS

Writing – review & editing: AND, XG, IS, PRM, SEQ, OP, BK, HSM

## SUPPORTING INFORMATION

**S1 Table. Participant demographics.** 19 participants who have Parkinson’s disease (PD) underwent single-unit recording in the substantia nigra pars reticulata (SNr). 3 participants who have PD and 2 participants who have Dystonia underwent single-unit recording in the globus pallidus internus (GPi). M = male; F = female; W = White; H = Hispanic.

**S1 Figure. Ultimatum game behavior. (A)** Overall reaction times in the computer (blue) and human (orange) conditions. *n* = 21 blocks of gameplay with human avatars and 8 blocks of gameplay with the computer avatar. **(B)** Overall mood ratings in the computer (blue) and human (orange) conditions. *n* = 7 blocks of gameplay with human avatars and 6 blocks of gameplay with the computer avatar. Bars represent mean, and error bars represent standard error of the mean.

**S1 Dataset. Parameter estimates.** Parameter estimates for all 10 candidate models applied to all behavioral session.

**S2 Dataset. Norm adaptation model comparison results for individual participants.** Bayesian information criterion (BIC) scores for each norm adaptation model for individual participants. Minimum BIC scores for each individual participant in each condition (human or computer) are in bold.

**S2 Figure. Comparison of models to Q learning.** Comparison of norm adaptation models to Q learning by total Bayesian information criterion (BIC) score for (**A**) all sessions, (**B**) human blocks only, and (**C**) computer blocks only. The model with the lowest total BIC score has the best fit (green). *n* = 29 sessions for (A), 21 sessions for (B), and 8 sessions for (C).

**S2 Table. Distributions of envy (*α*).** Distributions of empirical estimates for envy fit by each model in which envy is a free parameter. *n* = 29 behavioral datasets.

**S3 Table. Model frequencies and protected exceedance probabilities.** ΔBIC represents difference in total BIC relative to the lowest-scoring model. Expected model frequency represents the posterior estimate of the proportion of participants best characterized by each model. Protected exceedance probability represents the probability that a model is more frequent than other candidate models, corrected for the possibility that differences in model frequencies arose by chance (Bayesian omnibus risk = 0.023) and performed at the participant level. *n* = 29 behavioral datasets.

**S4 Table. Individual model recovery probabilities.** Model recovery probabilities for all 10 norm adaptation models. 29 synthetic behavioral datasets were simulated using parameters sampled from normal distributions based on the empirical mean and standard deviation for each parameter of each model fit to the 29 empirical behavioral datasets in our study. All 10 models were then refit to these simulated datasets. This process was repeated for 200 independent repetitions, with newly sampled parameter values and simulated datasets generated for all 29 synthetic subjects in each repetition. *n* = 29 simulated sessions repeated for 200 independent repetitions.

**S3 Figure. Individual model recovery matrix.** Model recovery matrix for all 10 norm adaptation models. *n* = 29 simulated sessions repeated for 200 independent repetitions.

**S5 Table. Class-level model recovery probabilities.** Class-level recovery probabilities generated by averaging recovery probabilities across models within each class (RW models, Bayes models, FS models, and the Q-learning model). *n* = 29 simulated sessions repeated for 200 independent repetitions.

**S4 Figure. Class-level model recovery matrix.** Class-level recovery matrix for the four model classes (RW models, Bayes models, FS models, and the Q-learning model). *n* = 29 simulated sessions repeated for 200 independent repetitions.

**S5 Figure. Parameter recovery.** Parameter recovery for **(A)** temperature, **(B)** initial norm, and **(C)** norm adaptation rate based on true (empirical) parameter estimates obtained by fitting models to 29 behavioral sessions and estimated (simulated) parameter estimates obtained by fitting models to 29 simulated behavioral sessions. Human and computer blocks were fitted separately. Behavior was fit with the RW norm adaptation model with variable initial norm and fixed alpha. \*\**p* < 0.01 and \*\*\**p* < 0.001 by Pearson’s product-moment correlation tests. *n* = 29 true (empirical) estimates and 29 estimated (simulated) estimates.

**S6 Figure. Posterior predictive checks.** Empirical vs. simulated behavior generated by the RW: fixed alpha model, including group-level (**A**) overall rejection rate by condition and (**B**) rejection rate by offer size in each condition. (**C**) Participant-level rejection rates for observed versus simulated sessions in the human and computer conditions. Human = orange, computer = blue. *n* = 29 empirical behavioral datasests and 29 simulated behavioral datasets.

**S7 Figure. Norm adaptation modeling for one example participant. (A)** Trial-by-trial choices, **(B)** internal norm estimates, and **(C)** norm prediction error (nPE) estimates for one example participant. Behavior was fit with the RW norm adaptation model with variable initial norm and fixed alpha.

**S6 Table. Patient-specific stereotactic coordinates of the microelectrode recordings collected for SNr putative DA neurons.** Patient-specific Leksell coordinates in X, Y, and Z space for each microelectrode recording of putative DA neuron activity in the SNr. Coordinates are in an AC-PC oriented space registered to the Schaltenbrand-Wahren atlas and each participant’s MRI and CT scans.

**S7 Table. Waveforms widths and average firing rates of SNr putative DA neurons.** Waveform width (ms) and average firing rate (Hz) for each putative DA neuron recorded in the SN.

**S8 Table. Summary of number of SNr putative DA neurons and task conditions played per recording session.** Summary of number of SNr putative DA neurons recorded and task conditions (human avatar or computer) played per participant per recording session. This summmary includes all recording sessions collected, including those later excluded from behavioral or neural analyses according to inclusion criteria.

**S8 Figure. Activity of one example SNr putative DA unit at split reveal. (A)** Raster, **(B)** peristimulus time histogram, and **(C)** smoothed continuous firing rate for one example putative DA unit at split reveal in the human condition. Split reveal occurs at time = 0 ms.

**S9 Table. Summary of mixed-effects model predicting SNr putative DA unit activity at split reveal.** Summary of mixed-effects model in which fixed effects nPE magnitude, nPE valence, condition (human avatar or computer) and their respective interaction terms with a random effect for unit predicted group activity of SNr putative DA units during the 1000 ms after split reveal. \*\**p* < 0.01 and *\*p* < 0.05 by mixed-effects model. Std. Error = standard error. *n* = 31 putative DA neurons.

**S9 Figure. Ultimatum game behavior and putative GPi neuron activity. (A)** Intraoperative targeting of the microelectrode in the GPi. **(B)** Rejection rate by offer size in the computer (blue) and human (orange) conditions for participants who played UG while undergoing single-unit recording in the GPi. **(C)** Average waveform for all GPi putative units (black) and average waveforms for individual GPi putative units (grey). **(D)** Probability density function of mean firing rates of GPi putative units during the 1000 ms after split reveal for trials producing positive (green) versus negative (magenta) nPEs in the human condition. Bars and lines represent the mean. Error bars represent standard error of the mean. Panel A created with BioRender.com. *n* = 6 blocks of gameplay with human avatars and 2 blocks of gameplay with the computer avatar for (B). *n* = 19 putative neurons for (C). n = 16 putative neurons in the human condition for (D). Brain slice created in BioRender. Neal, A. (2027) https://BioRender.com/8bs2hqw.

**S10 Table. Waveform widths and average firing rates of all GPi putative neurons.** Waveform width (ms) and average firing rate (Hz) for all putative neurons recorded in the GPi.

**S10 Figure. SNr putative DA unit activity before mood rating selection.** Mean firing rate of SN putative DA units during the 1000 ms before mood rating selection versus z-scored mood rating across both human and computer conditions. *n* = 12 putative DA neurons.

