## Supplementary material for "Single neurons in the human substantia nigra encode social learning signals": S1 Table

| **Recording target** | **Diagnosis** | **Age** | **Sex** | **Race** |
| --- | --- | --- | --- | --- |
| SNr | PD | 62 | M | W |
|  |  | 61 | M | W |
|  |  | 52 | M | W |
|  |  | 64 | M | W |
|  |  | 63 | M | W (H) |
|  |  | 66 | M | W |
|  |  | 76 | F | W |
|  |  | 75 | M | W |
|  |  | 67 | F | W |
|  |  | 64 | M | W |
|  |  | 74 | M | W |
|  |  | 68 | M | W |
|  |  | 66 | M | W |
|  |  | 65 | M | W |
|  |  | 68 | M | W |
|  |  | 65 | M | W |
|  |  | 39 | M | W |
|  |  | 73 | M | W |
|  |  | 67 | F | W |
| GPi | PD | 62 | M | W |
|  |  | 47 | M | W |
|  |  | 68 | M | W |
|  | Dystonia | 65 | M | W |
|  |  | 22 | M | W |
