## Supplementary material for "Single neurons in the human substantia nigra encode social learning signals": S2 Table

| **Model** | **Condition** | **Mean [IQR]** |
| --- | --- | --- |
| Bayes: fixed f0 | Computer | 0.89 [0.80–1.00] |
| Bayes: fixed f0 | Human | 0.90 [0.79–1.00] |
| Bayes: variable f0 | Computer | 1.00 [1.00–1.00] |
| Bayes: variable f0 | Human | 0.93 [1.00–1.00] |
| FS: fixed f0 | Computer | 0.90 [0.80–1.00] |
| FS: fixed f0 | Human | 0.90 [0.92–1.00] |
| FS: variable f0 | Computer | 0.61 [0.41–0.74] |
| FS: variable f0 | Human | 0.73 [0.70–0.80] |
| Q-learning | Computer | 0.61 [0.32–0.75] |
| Q-learning | Human | 0.46 [0.27–0.73] |
| RW: fixed f0 | Computer | 0.94 [0.88–1.00] |
| RW: fixed f0 | Human | 0.88 [0.78–1.00] |
| RW: variable f0 | Computer | 0.98 [0.96–1.00] |
| RW: variable f0 | Human | 0.92 [0.86–1.00] |
