## Supplementary material for "Single neurons in the human substantia nigra encode social learning signals": S3 Table

| **Model** | **ΔBIC** | **Expected Model Frequency** | **Protected Exceedance Probability** |
| --- | --- | --- | --- |
| RW: variable f0 | 30.138 | 0.062 | 0.004 |
| RW: fixed f0 | 188.583 | 0.046 | 0.003 |
| FS: fixed alpha | 0.000 | 0.435 | 0.932 |
| FS: variable f0 | 22.242 | 0.060 | 0.004 |
| FS: fixed f0 | 148.521 | 0.053 | 0.003 |
| Bayes: fixed alpha | 224.058 | 0.043 | 0.003 |
| Bayes: variable f0 | 240.451 | 0.043 | 0.003 |
| Bayes: fixed f0 | 268.512 | 0.043 | 0.003 |
| RW: fixed alpha | 13.707 | 0.174 | 0.043 |
| Q-learning | 406.564 | 0.042 | 0.003 |
