## Supplementary material for "Single neurons in the human substantia nigra encode social learning signals": S4 Table

|  | **Recovered model** | | | | | | | | | | |
| --- | --- | --- | --- | --- | --- | --- | --- | --- | --- | --- | --- |
| **Generating model** |  | RW: fixed alpha | RW: variable f0 | RW: fixed f0 | Bayes: fixed alpha | Bayes: variable f0 | Bayes: fixed f0 | FS: fixed alpha | FS: variable f0 | FS: fixed f0 | Q-learning |
|  | RW: fixed alpha | 0.4547 | 0.0079 | 0.0203 | 0.2738 | 0.0193 | 0.0610 | 0.1205 | 0.0016 | 0.0407 | 0.0002 |
|  | RW: variable f0 | 0.4576 | 0.0128 | 0.0186 | 0.2691 | 0.0131 | 0.0600 | 0.1243 | 0.0014 | 0.0428 | 0.0003 |
|  | RW: fixed f0 | 0.1338 | 0.0143 | 0.0972 | 0.2341 | 0.0160 | 0.3128 | 0.1172 | 0.0000 | 0.0740 | 0.0005 |
|  | Bayes: fixed alpha | 0.1641 | 0.0047 | 0.0017 | 0.4588 | 0.0029 | 0.0436 | 0.2003 | 0.0002 | 0.1157 | 0.0079 |
|  | Bayes: variable f0 | 0.1490 | 0.0048 | 0.0045 | 0.4502 | 0.0041 | 0.0569 | 0.1986 | 0.0002 | 0.1252 | 0.0066 |
|  | Bayes: fixed f0 | 0.0836 | 0.0060 | 0.0050 | 0.3155 | 0.0047 | 0.0933 | 0.2805 | 0.0000 | 0.2003 | 0.0110 |
|  | FS: fixed alpha | 0.0284 | 0.0022 | 0.0310 | 0.1478 | 0.0028 | 0.0921 | 0.3762 | 0.0000 | 0.3191 | 0.0003 |
|  | FS: variable f0 | 0.0860 | 0.0002 | 0.0078 | 0.0162 | 0.0009 | 0.0064 | 0.7200 | 0.0071 | 0.1550 | 0.0005 |
|  | FS: fixed f0 | 0.0683 | 0.0002 | 0.0222 | 0.0262 | 0.0000 | 0.0050 | 0.4188 | 0.0000 | 0.4583 | 0.0010 |
|  | Q-learning | 0.0128 | 0.0000 | 0.0000 | 0.1134 | 0.0000 | 0.0093 | 0.3110 | 0.0000 | 0.0455 | 0.5079 |
