## Supplementary material for "Single neurons in the human substantia nigra encode social learning signals": S5 Table

|  | **Recovered model** | | | | |
| --- | --- | --- | --- | --- | --- |
| **Generating model** |  | RW | Bayes | FS | Q-learning |
|  | RW | 0.4057 | 0.4198 | 0.1741 | 0.0003 |
|  | Bayes | 0.1411 | 0.4767 | 0.3737 | 0.0085 |
|  | FS | 0.0821 | 0.0991 | 0.8182 | 0.0006 |
|  | Q-learning | 0.0128 | 0.1228 | 0.3566 | 0.5079 |
