## Supplementary material for "Single neurons in the human substantia nigra encode social learning signals": S6 Table

| **Putative Neuron** | **X** | **Y** | **Z** |
| --- | --- | --- | --- |
| 92 | 87.6 | 101.9 | 89.2 |
| 460 | 87.6 | 101.9 | 89.2 |
| 651 | 107.5 | 100.2 | 90.4 |
| 667 | 85.7 | 97.9 | 85.9 |
| 414 | 106.8 | 92.9 | 93.1 |
| 369 | 110.6 | 99.9 | 90 |
| 412 | 110.6 | 99.9 | 90 |
| 490 | 87.8 | 101.6 | 85.8 |
| 229 | 90.4 | 105.9 | 92.4 |
| 252 | 90.4 | 105.9 | 92.4 |
| 364 | 87.7 | 107 | 100.4 |
| 373 | 87.7 | 107 | 100.4 |
| 354 | 107.5 | 110.9 | 119.4 |
| 383 | 107.5 | 110.9 | 119.4 |
| 224 | 87.6 | 102.6 | 96.9 |
| 245 | 87.6 | 102.6 | 96.9 |
| 996 | 87.2 | 99.7 | 87.5 |
| 1357 | 87.2 | 99.7 | 87.5 |
| 385 | 90.7 | 109.2 | 94 |
| 389 | 90.7 | 109.2 | 94 |
| 670 | 110.1 | 104.4 | 97.4 |
| 692 | 110.1 | 104.4 | 97.4 |
| 693 | 110.1 | 104.4 | 97.4 |
| 719 | 89.2 | 103.7 | 85.5 |
| 859 | 89.2 | 103.7 | 85.5 |
| 878 | 89.2 | 103.7 | 85.5 |
| 87 | 111.1 | 98.6 | 82.4 |
| 216 | 111.1 | 98.6 | 82.4 |
| 342 | 105.5 | 94 | 86.6 |
| 775 | 105.5 | 94 | 86.6 |
| 453 | 106.3 | 98.1 | 82.7 |
| 516 | 106.3 | 98.1 | 82.7 |
| 1208 | 110 | 103.7 | 99 |
| 1254 | 110 | 103.7 | 99 |
| 723 | 85.9 | 101.1 | 88.9 |
| 817 | 85.9 | 101.1 | 88.9 |
| 690 | 108.5 | 95.3 | 84.1 |
| 705 | 108.5 | 95.3 | 84.1 |
| 28 | 88.7 | 98.6 | 85.8 |
| 857 | 88.7 | 98.6 | 85.8 |
