## Supplementary material for "Single neurons in the human substantia nigra encode social learning signals": S7 Table

| **Putative neuron** | **Waveform width (ms)** | **Average firing rate (Hz)** |
| --- | --- | --- |
| 92 | 1.69 | 10.70 |
| 460 | 1.88 | 1.24 |
| 651 | 1.4 | 0.64 |
| 667 | 1.46 | 1.50 |
| 414 | 1.74 | 0.30 |
| 369 | 2.41 | 0.78 |
| 412 | 2.26 | 9.25 |
| 490 | 1.75 | 0.35 |
| 229 | 1.19 | 1.27 |
| 252 | 1.93 | 5.44 |
| 364 | 2.22 | 8.29 |
| 373 | 2.24 | 0.84 |
| 354 | 1.81 | 0.37 |
| 383 | 1.85 | 8.10 |
| 224 | 1.81 | 2.58 |
| 245 | 1.32 | 0.49 |
| 996 | 1.72 | 0.29 |
| 1357 | 1.72 | 14.63 |
| 385 | 1.71 | 2.09 |
| 389 | 1.77 | 1.97 |
| 670 | 1.89 | 0.21 |
| 692 | 1.86 | 0.79 |
| 693 | 1.69 | 7.86 |
| 719 | 1.99 | 1.36 |
| 859 | 1.52 | 1.00 |
| 878 | 1.46 | 6.98 |
| 87 | 1.78 | 0.34 |
| 216 | 1.79 | 11.06 |
| 342 | 1.18 | 0.14 |
| 775 | 1.63 | 15.24 |
| 453 | 1.91 | 0.53 |
| 516 | 1.93 | 7.58 |
| 1208 | 1.69 | 7.69 |
| 1254 | 1.84 | 1.95 |
| 723 | 1.74 | 5.76 |
| 817 | 1.65 | 6.98 |
| 690 | 1.92 | 0.72 |
| 705 | 1.24 | 5.61 |
| 28 | 1.36 | 0.84 |
| 857 | 1.61 | 11.98 |
