## Supplementary material for "Single neurons in the human substantia nigra encode social learning signals": S8 Table

| **Participant** | **Condition(s)** | **Number of Putative DA Neurons** |
| --- | --- | --- |
| pt0264_2 | Human | 2 |
| pt0265 | Human | **1** |
| pt0265_2 | Human | 1 |
| pt0267 | Human | 1 |
| pt0270 | Human | 0 |
| pt0270_2 | Human | 0 |
| pt0271 | Human | 2 |
| pt0271_2 | Human | 1 |
| pt0272 | Human | 0 |
| pt0272_2 | Human | 2 |
| pt0343_2 | Human | 0 |
| pt0358_2 | Human | 2 |
| pt0360 | Human | 2 |
| pt0374 | Human | 0 |
| pt0374_2 | Human | 2 |
| pt0296_2 | Human, Computer | 2 |
| pt0297_2 | Human, Computer | 2 |
| pt0298 | Human, Computer | 3 |
| pt0298_2 | Human, Computer | 3 |
| pt0377 | Human | 2 |
| pt0377_2 | Human, Computer | 0 |
| pt0378_2 | Human, Computer | 2 |
| pt0382 | Human, Computer | 2 |
| pt0383 | Human, Computer | 2 |
| pt0406 | Human, Computer | 2 |
| pt0409 | Human, Computer | 2 |
| pt0409_2 | Human, Computer | 2 |
