## Supplementary material for "Single neurons in the human substantia nigra encode social learning signals": S9 Table

| **Fixed effect** | **Estimate** | **Std. error** | **t-value** | **p-value** |
| --- | --- | --- | --- | --- |
| Intercept | -0.008979 | 0.02311 | -0.389 | 0.69832 |
| nPE magnitude | -0.0009855 | 0.002899 | -0.340 | 0.73439 |
| **nPE valence **** | 0.2016 | 0.07274 | 2.771 | **0.00566** |
| Condition | 0.008821 | 0.03674 | 0.240 | 0.81031 |
| nPE magnitude x nPE valence | -0.07129 | 0.03897 | -1.829 | 0.06758 |
| nPE magnitude x condition | -0.002443 | 0.004986 | -0.490 | 0.62450 |
| **nPE valence x condition *** | -0.2230 | 0.1063 | -2.097 | **0.03616** |
| nPE magnitude x nPE valence x condition | 0.08176 | 0.04574 | 1.788 | 0.07407 |
