## Supplementary material for "Single neurons in the human substantia nigra encode social learning signals": S10 Table

| **Putative neuron** | **Waveform width (ms)** | **Average firing rate (Hz)** |
| --- | --- | --- |
| 306 | 1.53 | 18.06 |
| 161 | 2.28 | 0.32 |
| 217 | 2.12 | 5.35 |
| 224 | 1.05 | 1.46 |
| 452 | 1.76 | 6.29 |
| 617 | 1.94 | 1.13 |
| 687 | 2.50 | 0.14 |
| 180 | 1.64 | 7.92 |
| 191 | 1.74 | 6.84 |
| 195 | 1.67 | 33.85 |
| 283 | 1.23 | 11.75 |
| 473 | 0.91 | 0.25 |
| 483 | 1.61 | 15.18 |
| 639 | 1.91 | 0.20 |
| 1149 | 1.32 | 32.28 |
| 1179 | 0.72 | 2.79 |
| 1231 | 0.63 | 4.52 |
| 1235 | 1.27 | 11.18 |
| 1243 | 0.58 | 0.32 |
